# Experimental and Phylogenetic Evidence for Correlated Gene Expression Evolution between Dermal and Endometrial Fibroblasts: implications for the evolution of cancer malignancy

**DOI:** 10.1101/2023.07.08.548194

**Authors:** Anasuya Dighe, Jamie Maziarz, Arig Ibrahim-Hashim, Robert A. Gatenby, Kshitiz, Andre Levchenko, Günter P. Wagner

**Affiliations:** Department of Ecology and Evolutionary Biology, Yale University, New Haven CT; Systems Biology Institute, Yale University, West Haven CT; Moffitt Cancer Center, Tampa, FL; Biomedical Engineering, University of Connecticut, Farmington, CT; Department of Biomedical Engineering, Yale University, New Haven, CT; Department of Evolutionary Biology, University of Vienna, Vienna Austria

## Abstract

Changes in transcriptional gene expression is a dominant mode of evolution, mostly driven by mutations at cis-regulatory regions. Mutations can affect gene expression in multiple cell types if the same cis-regulatory elements are used by different cell types. As a consequence, changes in gene expression in one cell type may be associated with similar gene expression changes in another cell type. Correlated gene expression change can explain correlated character evolution, as for instance the correlation between placental invasion and vulnerability to cancer malignancy. Here we test this hypothesis using a comparative and an experimental data set. Specifically, we investigate gene expression in dermal skin fibroblasts (SF) and uterine endometrial stomal fibroblasts (ESF). The comparative dataset consists of transcriptomes from cultured SF and ESF from 9 mammalian species. We calculated the independent phylogenetic contrasts (PIC) for each gene and cell type. We find that evolutionary changes in gene expression in SF and ESF are highly correlated, supporting the hypothesis that the correlated gene expression changes are a prevalent feature of gene expression evolution. The experimental data set derives from a SCID mouse strain that was selected for slow cancer growth which led to substantial changes in the SF compared to wild type SCID mice. We isolated SF and ESF from wild type and evolved SCID mice and compared their gene expression profiles. We find a significant correlation between the gene expression contrasts of SF and ESF, which supports the hypothesis that gene expression variation in SF and ESF is correlated. We discuss the implications of these findings for the hypothesized correlation between placental invasiveness and vulnerability to metastatic cancer.

## Introduction

Cell types are the fundamental building blocks of multicellular organisms ^1,2^. In evolution cell types arise through the gradual divergence of the gene regulatory networks of different cell populations, eventually leading to distinct although overlapping gene expression profiles^3 4-6^. More closely related cell types are more likely to have similar gene regulatory networks than more distantly related cell types. Therefore, closely related cell types are likely to be affected by the same mutations leading to correlated patterns of genetic variation in gene expression^4^, and, as a consequence, are expected to evolve similar gene expression changes among species^7^. In quantitative genetics this phenomenon is called “correlated selection response” ^8,9^. An example are fibroblastic cells, which exist in many tissues with tissue specific identities like stellate cells in the liver, or endometrial stromal fibroblasts in the uterus. Nevertheless, fibroblasts share phenotype defining gene expression programs, such as those for remodeling the extracellular matrix (ECM). Mutational changes in this ECM modifying program have the potential to affect the phenotype of different fibroblastic cell types. Here we test this prediction with respect to gene expression in the dermal (skin) fibroblast (SF) and the uterine endometrial stromal fibroblast (ESF). Specifically, we test two complementary predictions: 1) that, in evolution, gene expression in SF and ESF tend to change together, in particular for ECM related genes; and 2) that selection on gene expression in one cell type, the SF, leads to similar gene expression changes in another cell type, the ESF.

We use two orthogonal data sets to test these predictions. For the first prediction we use a comparative transcriptomic dataset from 9 mammalian species from the Boroeutheria clade of eutherian mammals and perform phylogenetic tests of correlated gene expression evolution. To test the second prediction, we use the results of an artificial selection experiment on SCID mice where male mice were selected for lower rate of cancer growth in an induced cancer model^10^.

The focus on skin fibroblasts (SF) and endometrial stromal fibroblasts (ESF) is motivated by their role in explaining the correlation between the structure of the fetal-maternal interface and the vulnerability of a species to cancer malignancy^11-13^. Species, from Eutheria (placental mammals *sensu stricto*), with less invasive placenta have endometrial cells that are more resistant to invasion by trophoblast cells than that of species with invasive placenta^12,14,15^. And species with less invasive placenta tend to be less vulnerable to cancer metastasis than species with invasive placenta, for instance humans^16^. This model has been called ELI for “Evolved Levels of Invasibility,” where the term “invasibility” is the vulnerability of a tissue to invasion by some cell type^17^. This term is well established in ecology, where it describes the vulnerability of a habitat to be invaded by foreign species^18,19^. In the case of tissues, the invasive cell type can be a cancer, the extra villous trophoblast cells of the placenta or a leukocyte. In brief, the model claims that the nature of the fetal-maternal interface is determined by the invasibility of the uterine endometrium (reviewed in^16^) and that the evolution of endometrial invasibility by placental cells will lead to correlated changes in somatic fibroblasts, thus leading to a corresponding change in invasibility of the body by cancer cells. The correlation of invasibility phenotypes has been tested experimentally^12,20^, but the idea of correlated gene expression evolution among these cell types has not. Here we present two orthogonal datasets to assess the validity of this assumption, one phylogenetic and the other experimental.

The phylogenetic test assesses whether, in evolution, gene expression changes in one cell type are correlated with corresponding gene expression changes in the other cell type. For this we take advantage of transcriptomic data from cultured of SF and ESF from 9 mammalian species described previously^20-22^. To test for correlated gene expression evolution, we use phylogenetic independent contrasts (PIC) ^23^. PICs estimate the amount of evolutionary change along independent parts of the phylogenetic tree. We find a strong signal of correlated gene expression evolution supporting a core assumption of the ELI model of cancer malignancy evolution, namely that evolution of gene expression in the uterus is correlated with gene expression evolution in the other fibroblasts. The direction of dependency, from uterus cells to skin cells, is motivated by the large differences in placental phenotype among mammalian species, suggesting fast evolution of placental invasiveness^24^.

For the experimental approach we focus on a SCID mouse line that was produced by selecting mice for slow tumor growth of subcutaneous injected Lewis-Lung Cancer cells ^10^. In this experiment the authors achieved a significant delay in the size increase of the experimental tumor. The design of the experiment ensured that the slower tumor growth was caused by changes in the host organism rather than changes to the tumor cells. Since SCID mice have no adaptive immune system, the likely cell type responsible for the slower tumor growth are mesenchymal cells, in particular skin fibroblasts. In fact, mice with slower growing tumors have a much more fibrous tissue composition of the dermal layer of the skin due to changes in the skin fibroblasts. We thus infer that artificial selection for tumor growth changed the gene expression in skin fibroblasts (as confirmed by our transcriptomic data below) and ask whether there are correlated changes in the gene expression profile of endometrial stromal fibroblasts (ESF). We find that gene expression differences between ESF from wild type and evolved SCID mice are correlated with the evolved changes in SF.

## Results

### Phylogenetic test for correlated gene expression evolution

We analyzed gene expression in cultured SF and ESF from 9 mammalian species representing two sub-clades of the Boroeutheria, namely the Laurasiatheria represented by dog, cat, horse, cow and sheep, as well as the Euarchontoglires represented by human, rat, guinea pig and rabbit (Figure 1A). These animals differ greatly in their fetal-maternal interface, where all Euarchontoglires have a hemochorial placenta (most invasive) and the Laurasiatheria representing less invasive phenotypes, such as epitheliochorial placenta in sheep, cow and horse as well as endotheliochorial in carnivores ^25^. In this taxon sample we expect that the ESF is the cell type under the selection, driven by evolution of different placental phenotypes, and changes in SF are mostly caused by correlated selection effects, since the function of the skin does not seem to be related to placental phenotype. This hypothesis can be assessed by observing the estimated amounts of evolutionary change in ESF compared to SF, where ESF are expected to display larger evolutionary changes than SF. This prediction is based on the quantitative genetic theory of correlated selection response^8^. This theory says that if two quantitative traits, say X and Y (e.g. expression of a gene in ESF=X and expression of the gene in SF=Y), are genetically correlated, any directional selection on one of these traits, say X, will lead to changes in the other character as well, say Y, even though Y is not itself under directional selection. The change in Y is called “correlated selection response.” However, the size of the correlated selection response in Y will be determined by the degree of correlation between X and Y, where lower correlations lead to smaller change in Y, assuming that X and Y have similar heritabilities.

**Figure 1:**
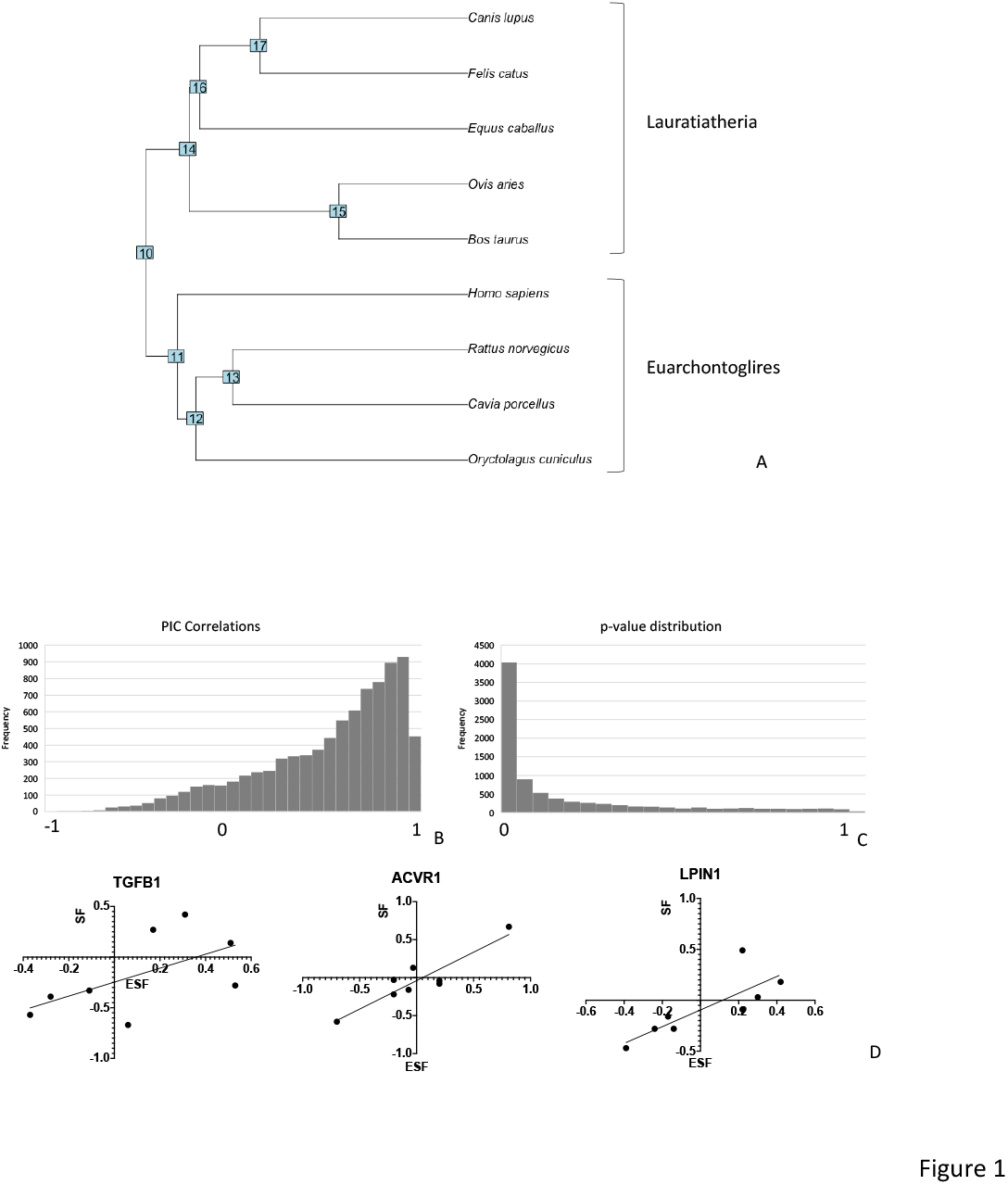
comparative test of correlated gene expression evolution between SF and ESF. A) phylogenetic relationships of species included in the comparison. B) distribution of correlation values among phylogenetic independent contrasts for gene expression changes in SF and ESF. C) distribution of p-values of the between PIC correlations showing strong bias towards small values. D) examples of genes with high PIC value correlations. The x-axis is always the PIC in ESF, the y-axis are the PIC values for SF.

We calculated the phylogenetic independent contrasts (PIC) for each gene in our taxon sample. Specifically, for each gene we calculated the 8 independent contrasts in both cell types, which estimate evolutionary gene expression changes in SF and ESF. For each gene we have a dataset of 8 PICs for SF and 8 corresponding PICs for ESF. We then calculated the correlation of the corresponding PICs for each gene. Figure 1B shows the distribution of correlation coefficients between corresponding PICs for ESF and SF for each of the one to one orthologs in this taxon sample. This distribution is heavily biased towards 1, with a mode within the interval [0.8, 0.9) representing 18% of the genes, and with 69% of genes with nominal correlations greater than 0.5. The distribution of p-values associated with these correlations (Figure 1C) allows us to estimate the number of genes that violate the null hypothesis of no correlation. About 80% of the 1-1 orthologous genes are predicted to have correlated gene expression evolution. Examples of a highly correlated genes in our data is TGFB1, ACVR1 and LPIN1 (Figure 1D). In order to assess whether contrasts in ESF are greater than that of SF we looked at the median slope is 0.66, indicating that the contrasts for SF tend to be smaller than that for ESF.

We calculated the GO enrichment of highly correlated genes for a strict criterion (p<0.001), and found that the corresponding FDR q-values of enriched GO categories is higher than 0.18, i.e. no strong enrichment of genes in any GO category (Suppl. Table 1). This result suggests that a broad set of genes are correlated in their evolutionary dynamic rather than genes from specific functional categories.

Overall, the pattern of gene expression divergence within the Boreoeutheria taxon sample is correlated between ESF and SF with a larger change in ESF than SF. This pattern is consistent with a model of correlated gene expression evolution in which gene expression in ESF is under directional selection and changes in SF are due to correlated selection responses.

While the comparative data analyzed above supports the notion that gene expression evolution in ESF and SF is correlated, we have not shown that gene expression is correlated because of genetic correlations. An alternative model would be that correlated gene expression evolution in these two cell types is due to co-selection on both cell types, i.e. that selection on gene expression for say ESF co-occurs with selection on gene expression in the other cell type, e.g. SF. In order to address this issue, we analyze a dataset where selection was only applied to one cell type and ask whether this leads to a correlated change in gene expression in the other cell type.

### Experimental test for correlated gene expression evolution

SCID mice from a strain selected for slow tumor growth^10^ have been obtained from the Charles River frozen embryo collection (MCC SCID line) and bred to form a colony at Yale University. We call these mice the SCID-EVOL strain. In addition, we obtained wild type SCID mice from from Charles River (Fox Chase SCID Beige Mouse, CB17.Cg-*Prkdc*^*scid*^*Lyst*^*bg-J*^/Crl) and established a colony that we call SCID-CTRL or SCID wild type. We first performed a transcriptomic analysis of skin fibroblasts isolated from the back skin. Data from four SCID-CTRL and three SCID-EVOL animals was obtained (described below). Next, we isolated endometrial stromal fibroblasts in order to test whether the uterus also exhibit gene expression changes due to correlated selection responses. To that end we monitored female SCID-CTRL and SCID-EVOL mice for their ovarian cycle stage via vaginal lavage and sacrificed animals when the vaginal lavage indicated the presence of large numbers of keratinized squamous epithelial cells, indicating estrus or metestrus I stage of the cycle. From the uterus ESF were isolated through differential adhesion and grown to near confluency before RNA extraction and transcriptome sequencing. QC analysis was performed on the expression profiles of protein coding nuclear genes that were expressed above the gene expression threshold of 3 TPM in at least one sample. The non-parametric Spearman correlation matrix (Figure 2A) shows that SF and ESF samples are well separated as expected for different cell types. The cluster analysis of the samples (Figure 2B) reveals that one of the control SF samples (SFCtrl2) clusters with the evolved samples for unknown reasons. We eliminated this sample from further analysis. Among the ESF samples, we identified one sample from an evolved animal to be strongly distinct in their gene expression profile from all the other samples (ESFEvol4) and two wild type samples (ESFCtrl1 and ESFCtrl2), which are different from other control samples. The latter two samples were from litter mates where one individual showed nasal secretions, suggesting that these two individuals may have been infected. All three outlier samples have been eliminated from further analysis, leaving seven ESF samples and six SF samples included in the analysis.

**Figure 2:**
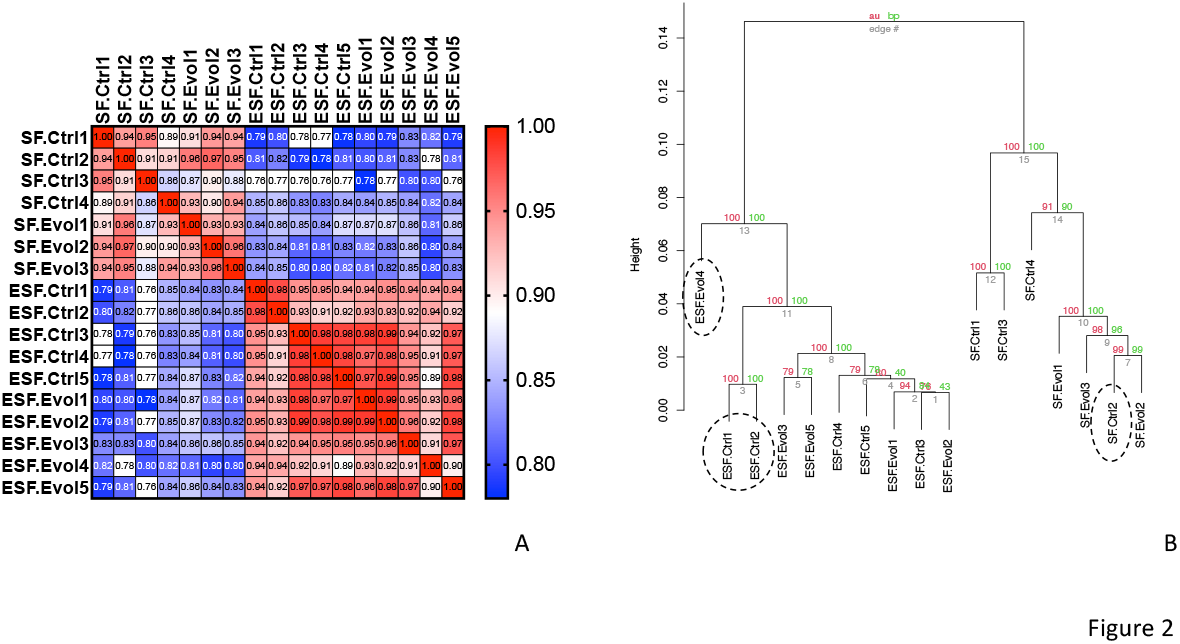
consistency analysis of all samples included in this study. A) heat map of Spearman correlation coefficients of the transcriptomic profiles; B) cluster analysis of samples identifying samples that were excluded from the analysis as indicated by dashed ovals. SF and ESF samples are well separated, as expected from different cell types. In the SF samples one Ctrl sample is nested within the evolved samples and has been eliminated. In the ESF samples one evolved and two controle samples are divergent from the rest and have been excluded from further analysis.

Figure 3A shows the distribution of the top 200 contrasts (=avTPM(Evol) - avTPM(Cntrl)) from SF samples. The highest contrast has been recorded at about 5400 TPM for the Thrombospondin 1 (*Thbs1*) gene, and the contrasts rapidly fall below 1000 TPM after 20 genes and approaches 100 TPM after the top 200 genes. The largest contrast in ESF is less than that for SF, at 2338 TPM for *Acta2*, Smooth muscle alpha2 actin, and also rapidly decreases to levels less than 500 TPM after the top 10 genes. Hence the average of the top contrasts for ESF is smaller (230TPM) than that of SF (475 TPM), as expected, if the change in ESF is due to correlated responses because of selection on gene expression in SF.

**Figure 3:**
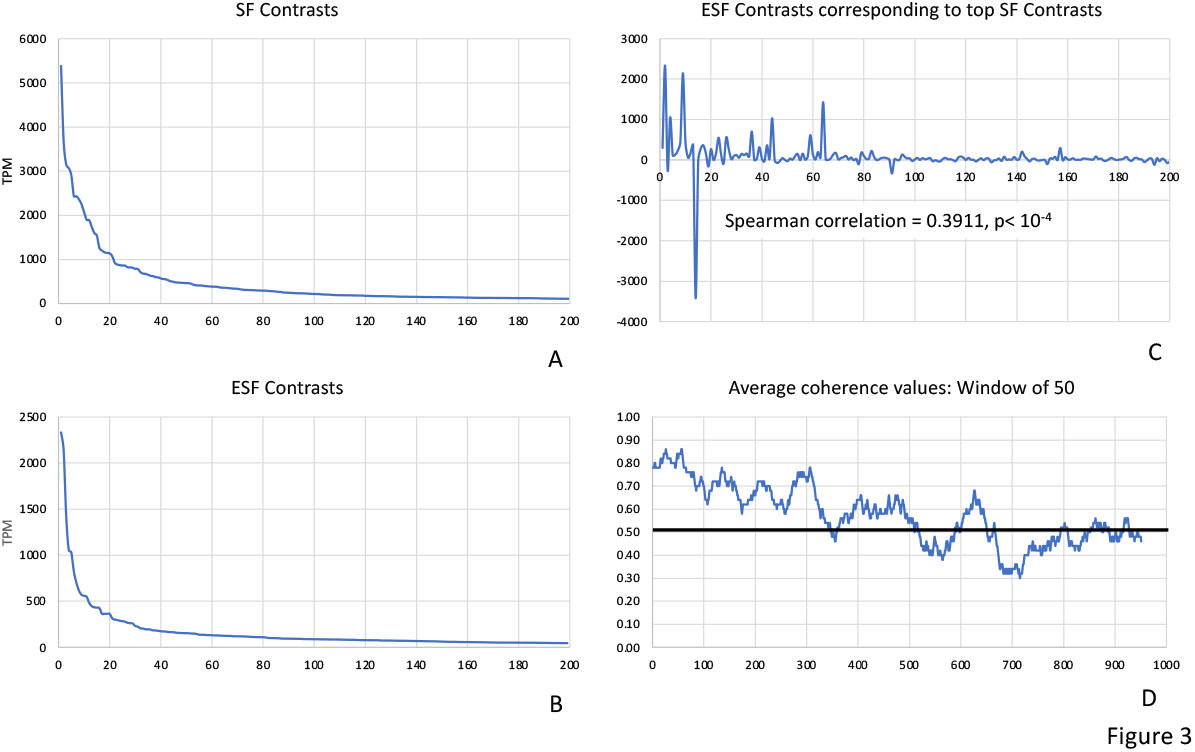
contrast profiles for different cell types. In each panel the x-axis is the order number of the genes sorted either by various criteria. A) Contrasts between evolved and control samples for SF samples. B) Contrasts between evolved and control samples for ESF samples. C) Contrasts between evolved and control samples for ESF samples where the genes are ordered by the size of SF contrasts (i.e. genes are arranged as in A). D) coherence statistic calculated over a sliding window of size 50. Random expectation of 0.5 is indicated by a thick black line.

The ESF contrasts of the genes that constitute the top 200 SF contrasts is highly variable, as expected (Figure 3C). To assess whether the ESF contrasts are more similar to SF contrasts than expected by chance we first calculated the non-parametric Spearman correlation coefficient and found it strongly positive (ρ=0.391, p<10^−4^), consistent with a correlated selection response. To further assess whether there was in fact a correlated selection response we calculated the relative frequency of contrasts that have the same sign in ESF as they have in SF for the top 1000 SF contrasts. We call this measure ‘coherence’ and calculate the coherence value over a sliding window of 50 genes (Figure 2D). The coherence measure starts between 0.8 and 0.9 among the genes with the highest SF contrast and slowly decrease to the random expectation of 0.5 after about 500 to 600 genes. The binomial test probability for coherence of 0.8 in 50 trials is p=1.19x10^−5^, also confirming the expectation of correlated gene expression evolution.

The genes with large gene expression contrast in SF and a congruent contrast in ESF are largely related to collagenous ECM and cytoskeleton. These are *Thbs1*, the collagen genes *Col1a1, Col1a2, Col3a1*, and Secreted Protein Acidic And Cysteine Rich, *Sparc*, as well as Biglycan, *Bgn* for collagenous ECM (Figure 4A, C). For cytoskeletal genes these include *Acta2*, Transglenin, *Tagln*, Vimentin, *Vim* (Figure 4B, D).

**Figure 4:**
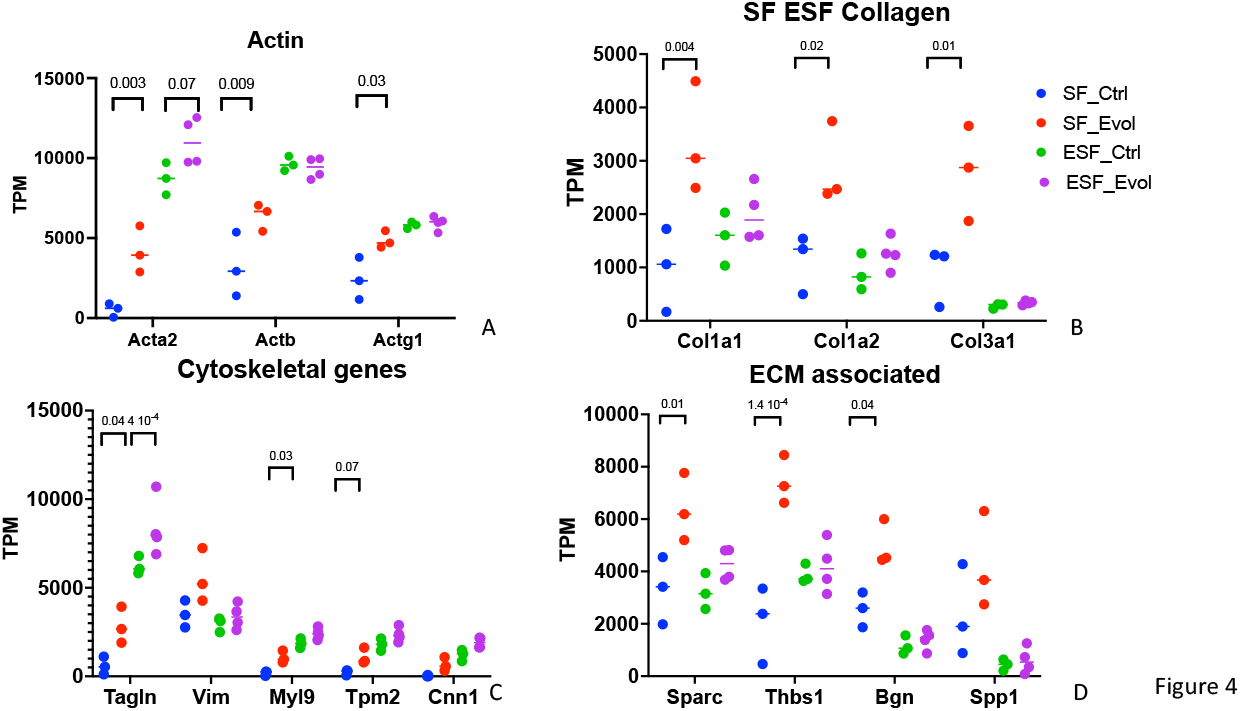
examples of genes with large SF contrast: A) actin genes, B) Collagen genes, C) other cytoskeletal genes, D) genes for ECM associated proteins.

These results support the hypothesis that the correlated evolution of gene expression in SF and ESF reported above can, at least in part, be due to genetic/mutational correlations in gene expression in these two cell types.

## Discussion

Because cell types originate in evolution by sister cell type divergence, they share parts of their gene regulatory network ^2,4,6,7^. One consequence of this model is that mutations that affect gene expression in one cell type tend to have, on average, some degree of correlated effects on gene expression in other cell types. If that is the case, quantitative genetic theory predicts that the evolution of gene expression in one cell type should lead to correlated gene expression changes in other cell types. We have tested this prediction here with a comparative phylogenetic and an experimental approach. In both cases the results are consistent with a model in which evolutionary changes in gene expression in one cell type is correlated with corresponding gene expression changes in a related cell type.

In the case of our comparative data, the species compared have been selected because they represent different phenotypes at the fetal-maternal interface. The taxon sample includes species with hemochorial placenta, i.e. species where the placenta erodes the maternal tissue and comes into direct contact with maternal blood, such a humans and rodents. On the other hand, there are species with none or only limited removal of maternal tissue as in the epitheliochorial and the endotheliochorial placenta found in cow, sheep and horse.

Experimental evidence suggests that the difference between invasive and non-invasive placentation is due to the endometrium, i.e. due to evolved differences in the maternal tissue. For instance, ectopic pregnancies in pigs lead to invasive implantation while in the uterus the placentation is non-invasive, i.e. epitheliochorial^14,15^. We also have shown in vitro that ESF from the cow are resisting invasion (i.e. are less invasible) by trophoblast cells, while human ESF are much more permissive to invasion, i.e. are more invasible^12^.

In the case of our experimental dataset the selection was on the growth rate of experimental tumors, which turned out to change skin fibroblasts. The involvement of skin fibroblasts is expected since SCID mice have no functioning adaptive immune system and the tumor cells were injected under the skin. At this site skin fibroblasts are forming the cancer associated stroma ^26^. There was no selection on ESF, since in the selective breeding was performed on males and males do not have ESF. Nevertheless, we found that genes which in SF show a large contrast between wild type and evolved populations also show a correlated response in ESF. The magnitude of the contrasts in SF is larger than that in ESF, which is consistent with the prediction that the changes in ESF are due to correlated selection responses to changes in SF rather than direct selection on ESF.

In both of our data sets the (natural or artificial) selection was on different types of fibroblasts. In the comparative dataset we assumed that species differences arose, in part, by selection on the endometrial stroma, i.e. the ESF, and the correlated response was on the skin fibroblasts. The rational was the fact that the maternal cells determine to a large degree the degree of placental invasion and the taxa samples differ from hemochorial to epitheliochorial placentation. This assumption is confirmed by the larger inferred evolutionary changes in ESF compared to that in SF. In the experimental dataset the artificial selection resulted in gene expression in SF, as shown by the large gene expression differences in SF while the ESF changes were smaller. As argued in the introduction, the cell phenotype determining features of fibroblasts are the expression of ECM molecules and that affecting cell shape (cytoskeletal genes). Consistently large differences in the expression of ECM and cytoskeletal genes are detected in ESF in the comparative data (suppl. Figure 1) and SF in the experimental data (Figure 4). The degree of expression correlation between cell types differs, as expected for two reasons. In the comparative data tens of million years of evolution has shaped the transcriptomes of these cells, with additional independent selection forces likely acting on both cell types. On the other hand, the selection response in the short term selection experiment depends on the spectrum of alleles coincidentally segregating in the founding population. Nevertheless, a broad consistency can be detected.

The results discussed in the previous paragraphs suggest that correlated gene expression evolution is pervasive and needs to be taken into account when studying gene expression evolution^7^. This fact is particularly important when considering the hypothesized correlation between cancer malignancy and the nature of the fetal-maternal interface^11^. Eutherian species with more invasive placenta tend to be more vulnerable to cancer malignancy, if tumor origination rate and malignancy rate are distinguished^13^. A number of genes have been shown to mediate this effect. One example is the evolution of the *CD44* gene which is expressed in much higher levels in humans and the rhesus monkey than in other eutherian species tested^21^. CD44 is a hyaluronic acid receptor and its expression in the tumor stroma has been implicated in cancer malignancy^27,28^. Similarly, we have shown that GATA2 is a pro-invasibility transcription factor, meaning that a knock out of *GATA2* in human SF and ESF leads to lower invasibility by trophoblast and cancer cells^20^. Similar results were obtained for *TFDP1*^*20*^. The finding that the evolution of gene expression in ESF and SF is correlated, and the fact that the same genes expressed in ESF and SF affect both placental and cancer invasion, supports a model where evolutionary changes at the fetal-maternal interface can affect the disease vulnerability of the respective species, in particular diseases with a similar cell biology as the embryo implantation such as cancer malignancy.

## Materials and Methods

### Mouse lines

SCID mice from a strain selected for slow tumor growth (Ibrahim-Hashim et al., 2020) were obtained from Charles River frozen embryo collection and bred to form a colony at Yale University (IACUC Protocol 2021-11483). We call these mice the SCID-EVOL strain. Wild type SCID mice were obtained from Charles River (Fox Chase SCID Beige Mouse, CB17.Cg-*Prkdc*^*scid*^*Lyst*^*bg-J*^/Crl). We call these mice SCID-CTRL.

### Ovarian cycle staging

Female SCID-CTRL and SCID-EVOL mice were monitored for their ovarian cycle stage via vaginal lavage. Lavage samples were spread onto glass slides and stained with hematoxylin and eosin. Animals were sacrificed when the vaginal lavage indicated the presence of large numbers of keratinized squamous epithelial cells, indicating estrus or metestrus I stage of the cycle.

### Isolation of human endometrial stromal cells and skin fibroblasts

An immortalized human ESF cell line was obtained from the Gil Mor group (ATCC-CRL-4003). Human SFs (BJ5ta) were purchased from ATCC (CRL-4001).

### Isolation of skin fibroblasts

Cow (Bos taurus), dog (Canis lupus), cat (Felis catus), guinea pig (Cavia porcellus), horse (Equus caballus), rabbit (Oryctolagus cuniculus), sheep (Ovis aries), rat (Rattus norvegicus), SCID-CTRL and SCID-EVOL SFs were obtained from fresh skin tissue as described previously^19^.

### Isolation of endometrial stromal fibroblasts

Cow (*Bos taurus*), dog (*Canis lupus*), cat (*Felis catus*), guinea pig (*Cavia porcellus*), horse (*Equus caballus*), rabbit (*Oryctolagus cuniculus*), sheep (*Ovis aries*), rat (*Rattus norvegicus*), SCID-CTRL and SCID-EVOL ESFs were obtained as described previously^19^.

### Cell culture

ESFs were grown in phenol-red free DMEM/F12 with high glucose (25 mM), supplemented with 10% charcoal-stripped calf serum (Gibco) and 1% antibiotic/antimycotic (Gibco). Human SFs BJ5ta (ATCC) cells were cultured in 80% DMEM and 20% MEM supplemented with 10% FBS, 1% antibiotic/antimycotic and 0.01 mg ml^−1^ hygromycin. All other SFs were cultured in DMEM with high glucose supplemented with 10% FBS.

### RNA isolation and sequencing

RNA was isolated using RNeasy micro kit (QIAGEN) and resuspended in 15 μl of water. The Yale Center for Genome Analysis ran samples on the Agilent Bioanalyzer 2100 to determine RNA quality, prepared mRNA libraries and sequenced on Illumina HiSeq2500 to generate 30–40 million reads per sample (single-end 75 base pair reads).

### Transcript-based abundances using RNAseq data

RNAseq data obtained were quantified using the transcript-based quantification approach as given in the program ‘kallisto’ ^29^. Here, reads are aligned to a reference transcriptome using a fast hashing of k-mers together with a directed de Bruijn graph of the transcriptome. This rapid quantification technique produces transcript-wise abundances which are then normalized and mapped to individual genes and ultimately reported in terms of TPM^30^. The Ensembl^31^ gene annotation model was used and raw sequence reads (single-end 75 bp) for ESFs and SFs from human (*Homo sapiens*), cow (*Bos taurus*), dog (*Canis lupus*), cat (*Felis catus*), guinea pig (*Cavia porcellus*), horse (*Equus caballus*), rabbit (*Oryctolagus cuniculus*), sheep (*Ovis aries*) and rat (*Mus musculus*) were aligned to GRCh38.p13, ARS-UCD1.2, CanFam3.1, Felis_catus_9.0, Cavpor3.0, EquCab3.0, OryCun2.0, Oar_v3.1 and Rnor_6.0 reference transcriptome assemblies. Further, to facilitate gene expression across species, a one-to-one ortholog dataset consisting of 8639 genes was formulated using the BioMart tool on Ensembl.

For quantifying gene expression from SCID RNAseq data (Ctrl and Evol), GRCm39 (GCA_000001635.9) mouse reference transcriptome from Ensembl was used.

### Calculation of Phylogenetic Independent Contrasts (PIC)

Phylogenetically independent contrasts (PIC) ^23,32,33^is a method for removing phylogenetic information from a character dataset with an aim to reduce the phylogenetic interdependence of data points, which can violate the assumptions of many statistical tests. The most broadly used methods to account for phylogenetic structure is the method of PICs where observations on N species are transformed into N-1 contrasts (differences). The method estimates the most likely history of evolutionary change in variables such that the contrasts reflect independent evolutionary changes. PIC values were computed by R package APE^34^ (Analysis of Phylogenetics and Evolution) using the function “pic” where the gene expression values (in TPM) were iteratively fed into the pic function to obtain an array of resultant PICs for each gene across 8 internal nodes in the phylogenetic tree used in this study.

**Supplemental Figure 1:**
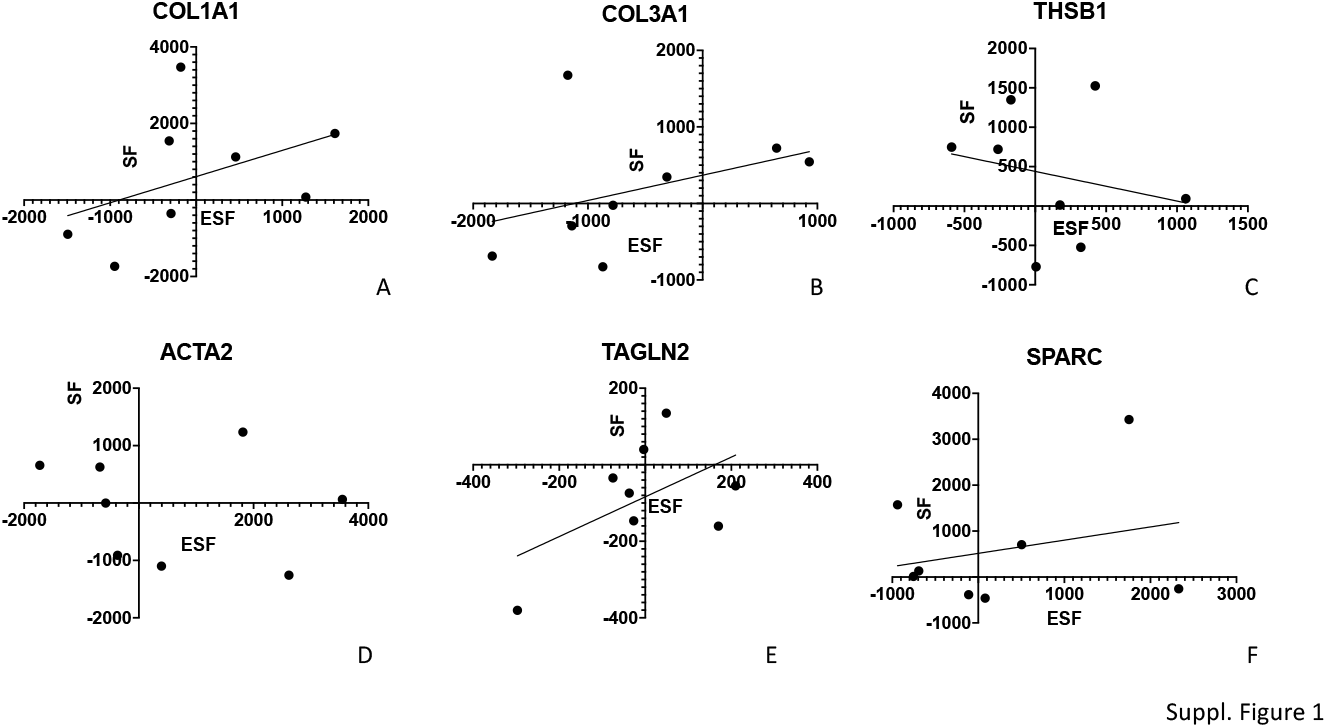
scattergrams of phylogenetic independent contrasts in ESF and SF for a number of genes which show direct effects in SF of the experimental SCID mice. A & B) *Col1a1* and *Col3a1* show a strong phylogenetic correlation between ESF and SF in the same way as it shows in the experimental population (see Figure 4A); C) *Thsb1* shows a strong evolutionary change in ESF but there is no correlated response in SF detectable, may be caused by different selective constraints on SF than ESF; D) Similarly to *Thsb1, Acta2* shows a large range of phylogenetic contrasts in ESF but the evolutionary changes in SF are not correlated to ESF; E) *Tagln2* in evolution shows a weak positive relationship but a strong one in experimental population (Figure 4C); F) *Sparc* is a gene for an ECM associated protein which shows a weak association between cell types in evolution as well as the experiment (Figure 4D).

**Supplemental Table 1:** gene function ontology analysis of genes which have a highly correlated PIC values. Note that the FDR q-value is high for all categories, suggesting that the correlated evolution does not affect very specific functional categories of genes.

| GO Term | Description | P-value | FDR q-value |
| --- | --- | --- | --- |
| GO:0007186 | G protein-coupled receptor signaling pathway | 1.36E-05 | 1.80E-01 |
| GO:0007610 | behavior | 3.71E-05 | 2.46E-01 |
| GO:0050727 | regulation of inflammatory response | 4.07E-05 | 1.80E-01 |
| GO:0050728 | negative regulation of inflammatory response | 6.37E-05 | 2.11E-01 |
| GO:0007165 | signal transduction | 7.03E-05 | 1.86E-01 |
| GO:0032501 | multicellular organismal process | 1.97E-04 | 4.34E-01 |
| GO:0007166 | cell surface receptor signaling pathway | 2.81E-04 | 5.32E-01 |
| GO:0032102 | negative regulation of response to external stimulus | 3.08E-04 | 5.10E-01 |
| GO:0003008 | system process | 3.58E-04 | 5.26E-01 |
| GO:0048609 | multicellular organismal reproductive process | 3.74E-04 | 4.96E-01 |
| GO:0006811 | ion transport | 4.90E-04 | 5.90E-01 |
| GO:0031348 | negative regulation of defense response | 4.95E-04 | 5.46E-01 |
| GO:0032101 | regulation of response to external stimulus | 5.40E-04 | 5.50E-01 |
| GO:0051239 | regulation of multicellular organismal process | 5.92E-04 | 5.60E-01 |
| GO:0098916 | anterograde trans-synaptic signaling | 6.35E-04 | 5.61E-01 |
| GO:0007268 | chemical synaptic transmission | 6.35E-04 | 5.26E-01 |
| GO:0007626 | locomotory behavior | 6.53E-04 | 5.09E-01 |
| GO:0050877 | nervous system process | 7.33E-04 | 5.40E-01 |
| GO:0060840 | artery development | 8.13E-04 | 5.67E-01 |
| GO:0032502 | developmental process | 9.43E-04 | 6.24E-01 |

